# Time series models for prediction of Leptospirosis in different climate zones in Sri Lanka

**DOI:** 10.1101/2021.02.19.431948

**Authors:** Janith Warnasekara, SB Agampodi, R Abeynayake

## Abstract

In tropical countries such as Sri Lanka, where leptospirosis—a deadly disease with a high mortality rate—is endemic, prediction is required for public health planning and resource allocation. Routinely collected meteorological data may offer an effective means of making such predictions. This study included monthly leptospirosis and meteorological data from January 2007 to April 2019 from Sri Lanka. Factor analysis was used first with rainfall data to classify districts into meteorological zones. We used a seasonal autoregressive integrated moving average (SARIMA) model for univariate predictions and autoregressive distributed lag (ARDL) model for multivariable analysis of leptospirosis with monthly average rainfall, temperature, relative humidity (RH), solar radiation (SR) and the number of rainy days/month (RD). Districts were classified into wet (WZ) and dry (DZ) zones and highlands (HL) based on the factor analysis of rainfall data. The WZ had the highest leptospirosis incidence; there was no difference in the incidence between the DZ and HL. Leptospirosis was fluctuated positively with rainfall, RH and RD, whereas temperature and SR were fluctuated negatively. ARIMA(1,0,0)(0,1,1)_12_, ARIMA(1,0,0)(1,1,1)_12_, and ARIMA(0,1,1)(0,1,1)_12_ were the best univariate models for DZ, WZ, and HL, respectively. Despite its known association, rainfall was positively significant in the WZ only at lag 5 (*P* = 0.03) but was negatively associated at lag 2 and 3 (*P* = 0.04). RD was positively associated for all three zones. Temperature was positively associated at lag 0 for the WZ and HL (*P* < 0.009) and was negatively associated at lag 1 for the WZ (*P* = 0.01). There was no association with RH in contrast to previous studies. Based on altitude and rainfall data, meteorological variables could effectively predict the incidence of leptospirosis with different models for different climatic zones. These predictive models could be effectively used in public health planning purposes.

## Introduction

Globally, leptospirosis accounts for nearly 1 million patients, 58,000 deaths and the loss of 2.9 million disability-adjusted life years (DALYs) annually(1,2). This disease, which is mainly associated with urban slum living conditions and occupational or recreational activities(3,4), is caused by a spirochete of the genus *Leptospira*(5). Any mammal, bird or marsupial can harbour and can have leptospirosis(6), although humans are considered an accidental host of *Leptospira*(5). *Leptospira* enter the body of the host animal via small abrasions or following contact with mucosal membranes. Before entering into the host, *Leptospira* must survive in water or in soil under different environmental conditions. Although there are free-living non-pathogenic saprophytic *Leptospira* in the environment, the survival of pathogenic *Leptospira* in the environment is not well understood. Ecological conditions conducive to *Leptospira* need to be evaluated to determine proper preventive measures for leptospirosis(7).

Climatic conditions affect the growth and survival of many microorganisms, and temperature, precipitation, relative humidity and wind play a role in vector-borne diseases(8). Rainfall, temperature, daily sunshine, relative humidity and solar radiation are associated with leptospirosis in many different settings(9–13). Moisture facilitates *Leptospira* growth, and thus an increase in the number of rainy days increases the survival of these bacteria. The optimal temperature for growth and survival for *Leptospira* is 30 °C. Although an optimal ambient temperature facilitates the growth and survival of *Leptospira*(14), high temperatures and increased solar radiation have a negative impact on rainfall and limit the survival of the organism. Relative humidity is also associated with disease occurrence(11,12). In addition, hot and humid conditions promote the growth and survival of rodents, which are a known vector for *Leptospira*(15–17). As environmental predictors are not uniform across different regions(13,15), research is needed to create regionally specific models to predict leptospirosis.

Sri Lanka is a hotspot for leptospirosis and has had several recent outbreaks(1,2,18,19), yet few studies have been carried out in Sri Lanka to determine associations between meteorological data and leptospirosis. A wavelet time series modelling done for the Kandy district showed associations of rainfall, the number of wet days per week, days with rainfall >100 mm per week, minimum temperature, average temperature and average humidity with leptospirosis(20). Two studies done by S. R. Gnanapragasam developed univariate seasonal autoregressive integrated moving average (SARIMA) models to describe the incidence of leptospirosis in Kurunegala, Rathnapura, Colombo, Gampaha and Kaluthara districts and for the western province.(21,22) A spatial epidemiological analysis by Robertson et al.(23) is the only study analyzing leptospirosis data from the dry zone of Sri Lanka, although meteorological parameters were not analysed. Plouffe Cameron et al. performed a comprehensive analysis to predict leptospirosis in Sri Lanka, but the only meteorological variable considered was rainfall and the major results presented were limited to the Colombo, Kaluthara and Mathale districts(24). These Sri Lankan studies were all focused on one area or one administrative district. However, epidemiological factors, clinical parameters and environmental conditions are different in the wet and dry zones of Sri Lanka(25,26). Also, the central highlands of the country behave dissimilarly to the dry and wet zones with respect to meteorological parameters(26). It is likely that the meteorological parameters associated with leptospirosis and/or the strength of their associations also differ across these climate zones. Therefore, the objective of this study was to determine the meteorological parameters associated with the incidence of leptospirosis and to evaluate the best leptospirosis prediction models for the wet and dry zones and highlands of Sri Lanka.

## Materials and Methods

### Data sources

We extracted disease incidence data publicly available through the disease surveillance system of Sri Lanka and obtained routinely collected meteorological data from the meteorology department of Sri Lanka(25,26). For the disease surveillance system, the first physician who suspects an incidence of the disease notifies the patient that he/she has a possible case of leptospirosis. This notification is done based on the surveillance case definition for notifiable diseases of Sri Lanka(27). Notifications are delivered via all government and private hospital/healthcare delivery institutions in Sri Lanka. Notification data are sent to the epidemiology unit, which is the national-level institution responsible for infectious disease control, through the medical officer of health divisions. The data summarized by district are published on a weekly basis(25). The leptospirosis notification process was streamlined as a result of a heavy outbreak reported during 2008(28). Therefore, we extracted weekly leptospirosis data from January 2007 until April 2019 (available at www.epid.gov.lk) and compiled monthly data separately for all 25 districts of Sri Lanka. District-level monthly meteorological data on rainfall, relative humidity, temperature, solar radiation and the number of rainy days per month from January 2007 to April 2019 were purchased from the meteorological department of Sri Lanka(26).

### Study setting

Although Sri Lanka is a small island, different geographical and climatic zones lead to diverse patterns of leptospirosis. There are several epidemiological and clinical differences in leptospirosis reported from different parts of Sri Lanka(23,29). There are three major climatic zones in Sri Lanka, namely the wet zone (annual rainfall, >2500 mm), intermediate zone (1750—2500 mm) and dry zone (<1750 mm) (Fig 1). The central highlands (hill country) of Sri Lanka overlap the dry, wet and intermediate zones and have low average temperatures, with an average minimum of 18 °C(26). Leptospirosis that occurs in the highlands is slightly different than that which occurs in the wet and dry zones, and thus the highlands were considered as a different geographical area for the purpose of this study(30). Cases of leptospirosis are commonly reported in the wet zone, whereas they are less frequently reported in the dry zone. The wet zone receives heavy rainfall associated with the southwestern monsoon rains, which favours the growth and survival of *Leptospira.* The dry zone environment is less favourable for *Leptospira*, as it receives rain only in association with northeastern monsoon rains, which occur from November to January(26). Several outbreaks of leptospirosis have, however, been reported in the dry zone since 2008(18,31).

**Fig 1.**
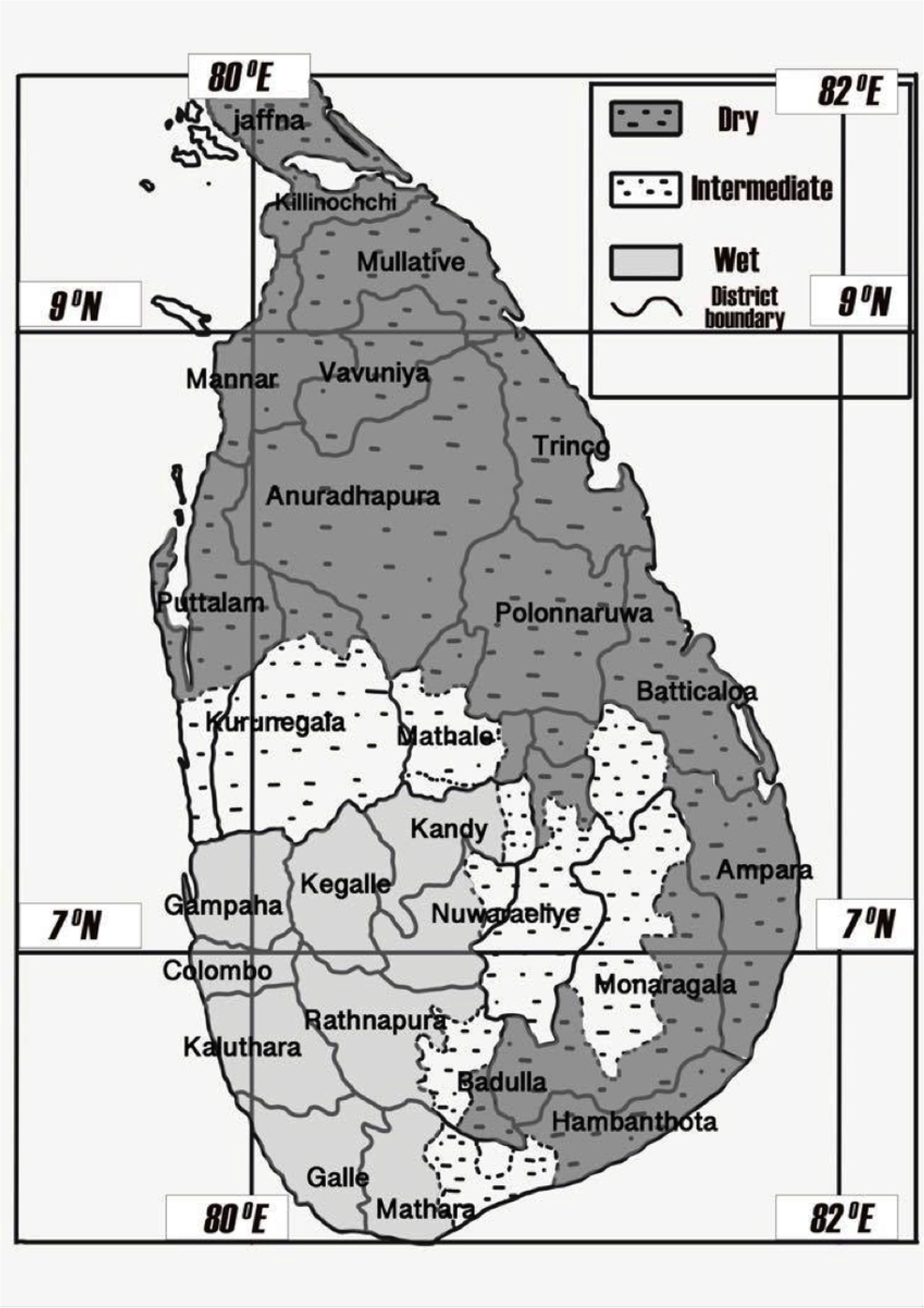
Distribution of climate zones and meteorological data collection centres in relation to administrative districts of Sri Lanka.

### Geographical area base analysis

For this study, Sri Lanka was classified into three different climatic zones based on rainfall. We used a factor analysis based on principal component factoring method for the rainfall data to confirm or redefine the traditional climatic zone categorization. Three districts, namely Badulla, Kurunegala and Hambanthota were excluded from the factor analysis as they each consisted of both dry and wet zones within their district boundaries. Table 1 shows the three geographical areas considered in the study and the districts belonging to each area.

**Table 1.** Districts and meteorological data collection sites within Sri Lanka according to meteorological zones.

| Dry Zone |  | Wet Zone |  | Highlands |  |
| --- | --- | --- | --- | --- | --- |
| Districts | Meteorological data collection centers | Districts | Meteorological data collection centers | Districts | Meteorological data collection centers |
| Jaffna | Jaffna | Colombo | Colombo | Kandy | Katugasthota |
| Mannar | Mannar | Gampaha | Katunayaka | Mathale | Nuwaraeliya |
| Kilinochchi | Vauniya | Kaluthara | Rathmalana | Nuwaraeliya |  |
| Vauniya | Anuradhapura | Galle | Galle |  |  |
| Mulathiv | Polonnaruwa | Mathara | Rathnapura |  |  |
| Anuradhapura | Puttalama | Rathnapura |  |  |  |
| Polonnaruwa | Tricomalee | Kegalle |  |  |  |
| Puttalama | Batticalo |  |  |  |  |
| Tricomalee | Monaragala |  |  |  |  |
| Batticalo | Potuvil |  |  |  |  |
| Ampara | Mahailluppallama |  |  |  |  |
| Monaragala |  |  |  |  |  |

### Data processing

The analysis was based on the refined meteorological zones as determined by the principal component factor analysis. For each area, we created the variable ‘monthly leptospirosis cases’ by summing all the reported cases of all the districts within that geographical area. Each variable was generated for 148 time points with data from January 2007 to April 2019. Then the average of each meteorological parameter was calculated for each area using the data of the districts within the area. Five meteorological variables were created for each zone using five meteorological parameters namely monthly rainfall, monthly rainy days, monthly relative humidity, monthly temperature and monthly solar radiation.

### Data analysis

#### Basic analysis

All data analysis was done using Statistical Package for Social Sciences (SPSS) version 23 and Eviews version 10. Summary statistics were obtained for all climatic and disease cases. The incidence of leptospirosis was calculated per 100,000 population in each zone and incidence numbers were compared using the Mann-Whitney *U*-test. To identify monthly variation in leptospirosis cases and meteorological parameters, the standardized (Z transformed) values of all variables were plotted in line diagrams. The Spearman correlation coefficient (*r*) was used as a measure of the association of leptospirosis cases with each meteorological parameter. Solar radiation was excluded from the multivariable analysis as a result of missing values.

#### Time series analysis

##### Models using previously reported leptospirosis data (univariate analysis)

SARIMA models were applied using SPSS to detect best-fitting univariate models for the three geographical zones. The model designation SARIMA/ARIMA (p, d, q) (P, D, Q)_s_ consists of regular autoregressive (AR-p) and moving average (MA-q) terms, which account for correlations with low lags. In addition, seasonal AR (P) and seasonal MA (Q) terms account for correlations with seasonal lags. For the purpose of this study, seasonality was analysed over a time frame of months i.e., there were 12 time periods in one season(Seasonality for the model is 12 months). The terms ‘D’ and ‘d’ indicate the number of seasonal differencing and regular differencing, respectively, used to make the mean of the data series stationary, and the term ‘s’ indicates the seasonality. The SARIMA model specification is {ARIMA (1,1,1) (1,1,1)_12_}

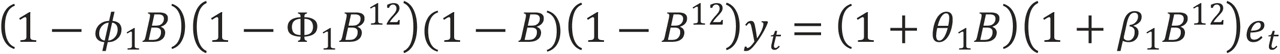

where (1 ― *ϕ*_1_*B*) = non-seasonal AR(1), (1 ― *Φ*_1_*B*^12^) = seasonal AR(1), (1 — B) = non-seasonal difference, (1 — B^12^) = seasonal difference, y_t_ = forecasted value, (1 + *θ*_1_*B*) = non-seasonal MA(1), (1 + *β*_1_*B*^12^) = seasonal MA(1), e_t_ = error term.

#### Model validation (univariate)

To detect the serial correlation of the univariate SARIMA models, the Ljung-Box Q statistic was used, and the Shapiro-Wilk test and Kolmogorov-Smirnov test were used to detect the normality of residuals. The Ljung-Box Q statistic was calculated using the following equation:

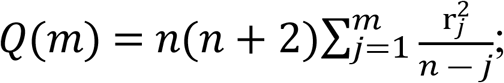

where Q(m) = Ljung-Box Q statistic, m = time lag, r_j_ = accumulated sample autocorrelation, n = length of the time series.

The test statistic (W) of the Shapiro-Wilk test was calculated as follows:

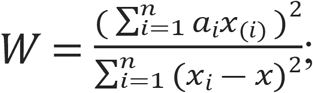

where x_(i)_ = i^th^ order statistics, a_i_ = coefficient calculated by the covariance matrix, *x* = sample mean.

The test statistic (D_n_) of the Kolmogorov-Smirnov test was calculated as follows:

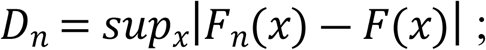

where sup_x_ = supremum of the set of distances and F = distribution function.

#### Models using climate variables with previously reported leptospirosis data (multivariable analysis)

##### Pretesting data

For multivariable analyses, all the variables were tested for mean stationarity using the unit root test of Eviews. In the unit root test, the stylized trend-cycle decomposition of a time series y_t_ is given by

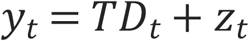

where TD_t_ is a deterministic linear trend and z_t_ is the AR(1) process.

In addition, outliers of the dependent variable were adjusted and interpolated using Eviews. Variance was stationary for all variables except the dependent variable (number of patients). Therefore, the dependent variable was transformed into a natural logarithm to make the data stationary for variance. Transformed data produced the best results in the zonal level analysis. Therefore, all climate zone level time series models presented in this paper include a natural log—transformed dependent variable.

#### Model fitting (multivariable)

The autoregressive distributed lag (ARDL) technique can be used for non-stationary data as well as for time series with a mixed order of integration. In our data set, three variables for dry zone data and two variables for wet zone data achieved a stationary mean in the 1^st^ difference{I(1)} (Table 6). There are several advantages of ARDL models. Irrespective of the difference order—I(0) or I(1) or a combination—ARDL models can be applied. Also, all underlying variables stand as a single equation in ARDL models. Therefore, endogeneity is less of a problem in ARDL models.(32)

The ARDL (p,q_1_,q_2_,q_3_,q_4_) model specification is

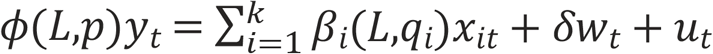

where

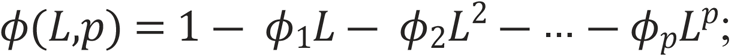

and

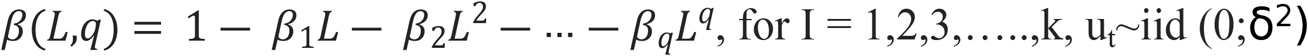

and where p = number of patients; q_1_, q_2_, q_3_ and q_4_ represent rainfall, rainy days, temperature and relative humidity, respectively; L = the lag operator and w_t_ = the vector of deterministic variables.

#### Model validation (multivariable)

Within the meteorological dataset the first 108 time points were used to create the models and the next 40 time points were used to validate the models. The lowest Akaike information criterion (AIC) and the lowest mean absolute percentage error (MAPE) were used for the selection of the best-fitting models:

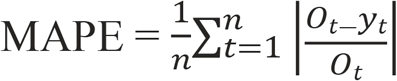

where n = number of time points, O_t_ = observed value, y_t_ = forecasted value and AIC = —(log-likelihood) + 2K

where **K** is the number of model parameters.

To identify serial autocorrelation of the best-[fitting] model, the Lagrange multiplier (LM) test was applied. The LM test is the product of the coefficient of determination (R^2^) from auxiliary regression and the sample size, LM = nR^2^, where R^2^ = coefficient of determination and n = number of time points. A non-significant (i.e., *P* **>**0.05) value for the LM test indicates that the residuals of the time series model are not serially auto correlated.

Residuals were checked for normality using the Jarque-Bera (JB) test, which is based on sample skewness and sample kurtosis. In addition, the residuals were tested for the presence of the ARCH effect to determine heteroscedasticity.

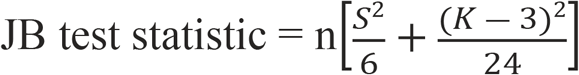

where S = skewness and K = kurtosis.

## Results

### Meteorological zone classification

The factor analysis extracted two components. The first component resulted in the grouping of meteorological data collection centres in the dry zone, and the second component resulted in the grouping of meteorological data collection centres in the wet zone. However, the Katugasthota (in the Kandy district) and Nuwaraeliya centres showed higher factor loadings for both components. These two districts are situated in the central highlands of the country. Therefore, these two districts and Mathale district (the only other district in the central highlands, which does not have a major meteorological centre) were considered as the “highlands”. Fig 2 shows the component matrix in rotated space.

**Fig 2.**
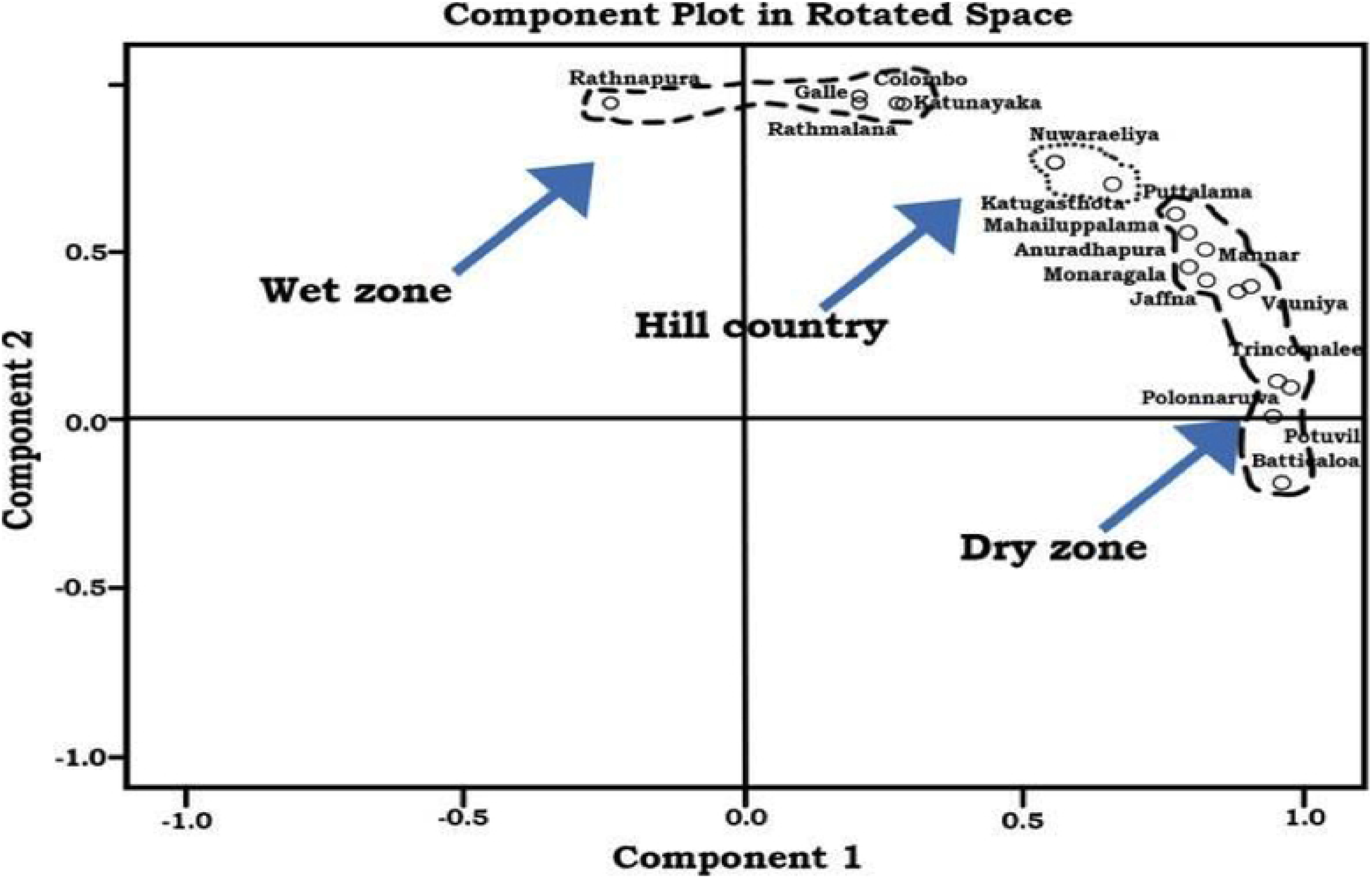
Classification of Sri Lankan climatic zones based on rainfall represented as a component plot in rotated space. The fig 2 clearly demarcate the dry areas towards component 1 (‘x’ axis) and wet areas towards component 2 (‘y’ axis) and highlands towards center. Accordingly, meteorological data collection centers and their respective districts were classified as belonging to the wet zone, dry zone or highlands (Table 1). Kurunegala, Badulla and Hambanthota districts were excluded from the initial categorization, as their geographical areas showed considerable overlap with these three climate zones.

### Descriptive analysis of zonal-level data

The incidence of leptospirosis was significantly higher in the wet zone as compared with the other two zones (Tables 2 and 3). The incidence was not significantly different between the dry zone and highlands, although the incidence was slightly higher in the highlands.

**Table 2.** Descriptive statistics of the model parameters used to describe the incidence of leptospirosis cases and meteorological variables.

| Parameter | Dry zone |  |  | Wet Zone |  |  | High Lands |  |  |
| --- | --- | --- | --- | --- | --- | --- | --- | --- | --- |
|  | Median | Min | Max | Median | Min | Max | Median | Min | Max |
| No of leptospirosis patients per month | 27.5 | 3 | 265 | 136.5 | 3 | 1023 | 18 | 0 | 219 |
| Monthly incidence of leptospirosis per 100,000 population | 0.54 | .06 | 5.2 | 1.42 | 0.03 | 10.59 | 0.70 | 0.00 | 8.52 |
| Rainfall | 79.5 | 1.6 | 559.3 | 187.1 | 7.6 | 744.8 | 131.2 | 3.6 | 554.4 |
| Relative Humidity | 79.0 | 70.2 | 88.8 | 82.8 | 75.4 | 87.3 | 85.0 | 68.5 | 91.6 |
| Temperature | 28.5 | 25.1 | 30.5 | 27.7 | 26.3 | 29.8 | 20.7 | 18.7 | 22.7 |
| Rainy days | 7.0 | 1.4 | 21.8 | 15.7 | 2.5 | 27.4 | 15.0 | 1.5 | 26.5 |

\*SD- Standard deviation; Min-minimum; Max-maximum; Md - Median

**Table 3.** Comparison of the number of leptospirosis cases among meteorological zones of Sri Lanka (Mann-Whitney *U*-test)

| Comparison | Incidence |  |
| --- | --- | --- |
|  | Z value | P value |
| Dry vs. Wet | 8.6 | 0.000 |
| Dry vs. High | 1.8 | 0.059 |
| Wet vs. High | 7.2 | 0.000 |

\* Z =z value; P = p value

### Monthly fluctuations in patient numbers in conjunction with meteorological data

Figs 3—5 show the monthly fluctuations of the standardized meteorological parameters with standardized patient numbers. In the dry zone, a single outbreak occurs from October to January, whereas two annual outbreaks occur in the wet zone (March and August to November) and highlands (May and October to January).

**Fig 3.**
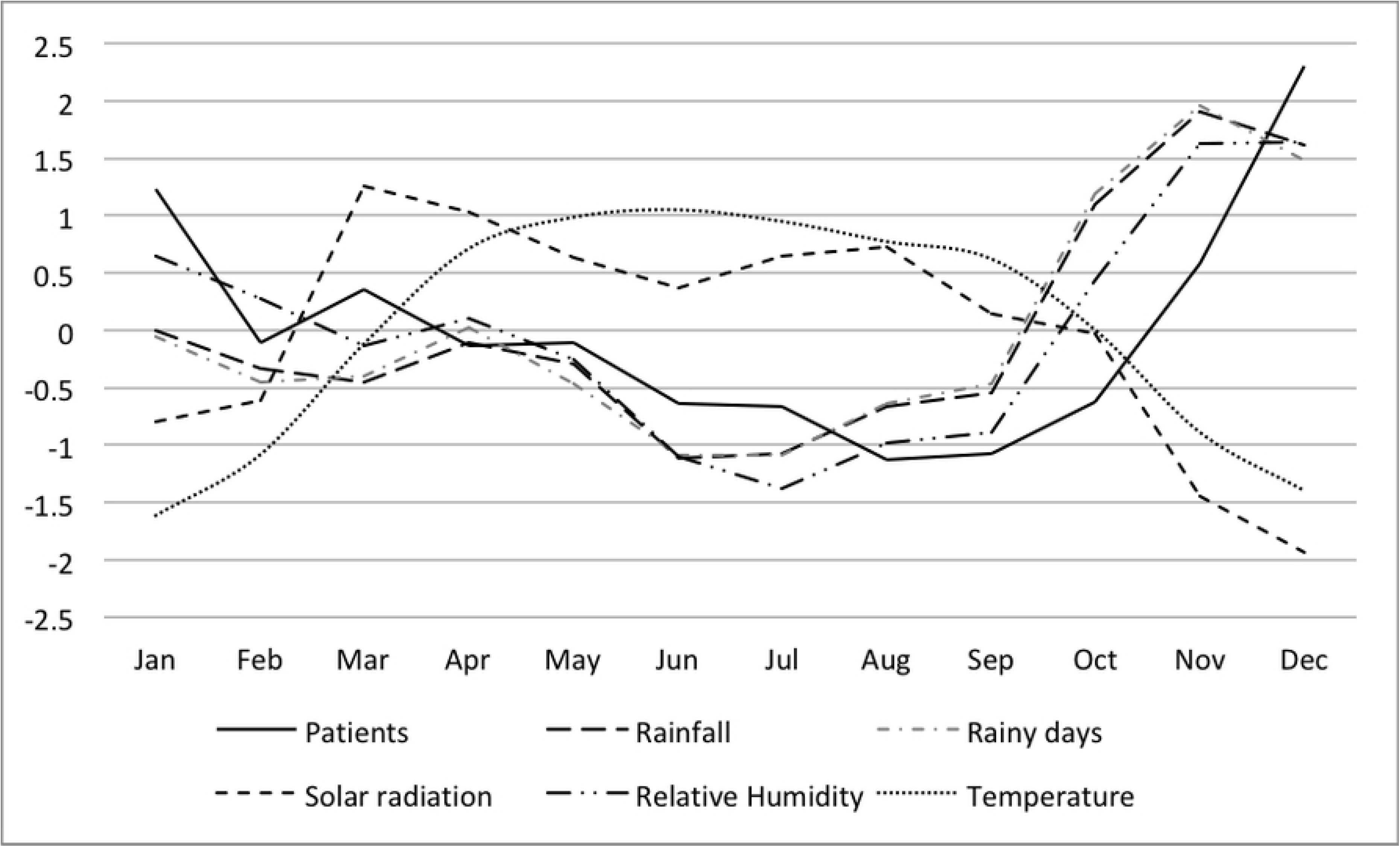
Monthly distribution of the standardized incidence of leptospirosis cases and meteorological parameters in the dry zone of Sri Lanka.

**Fig 4.**
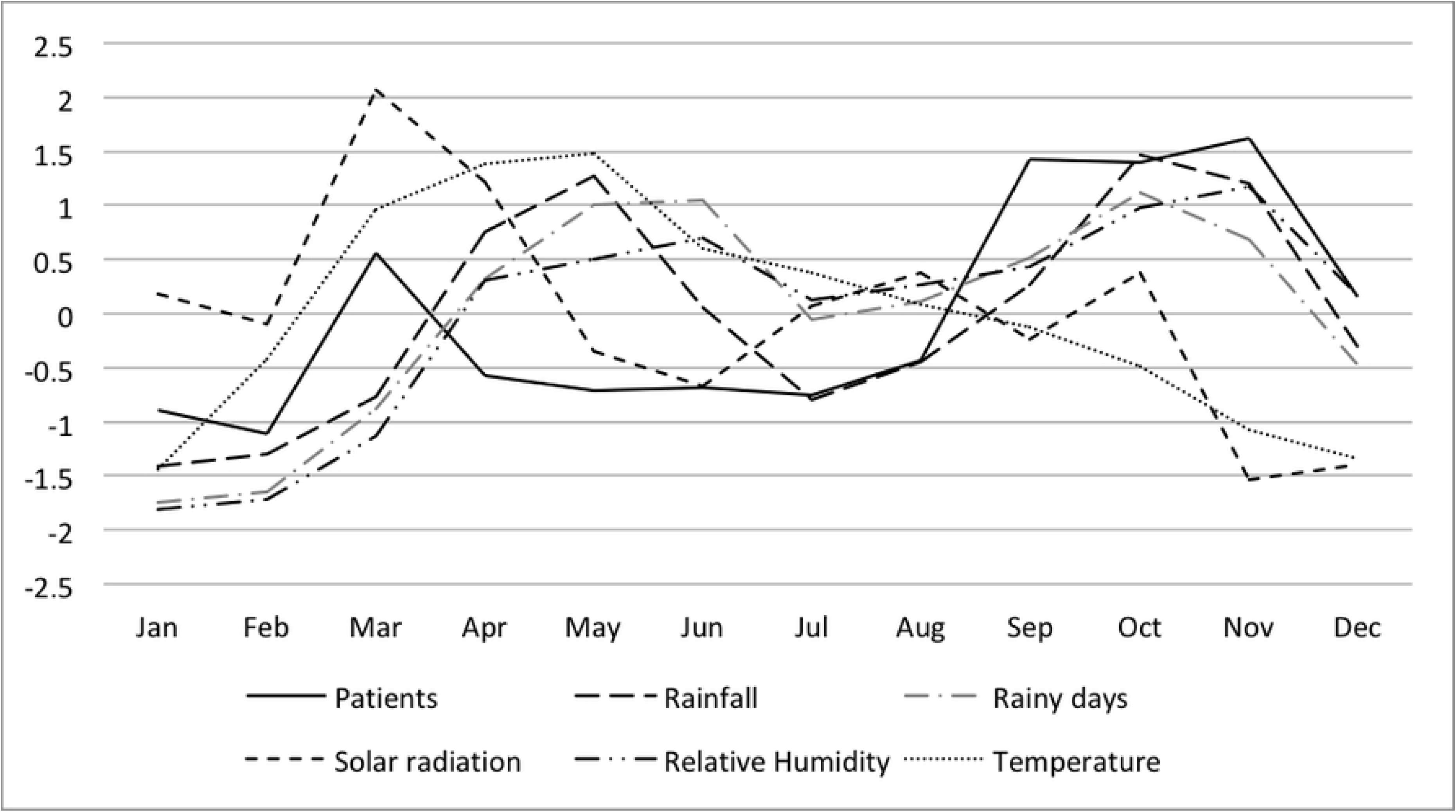
Monthly distribution of the standardized incidence of leptospirosis cases and meteorological parameters in the wet zone of Sri Lanka.

**Fig 5.**
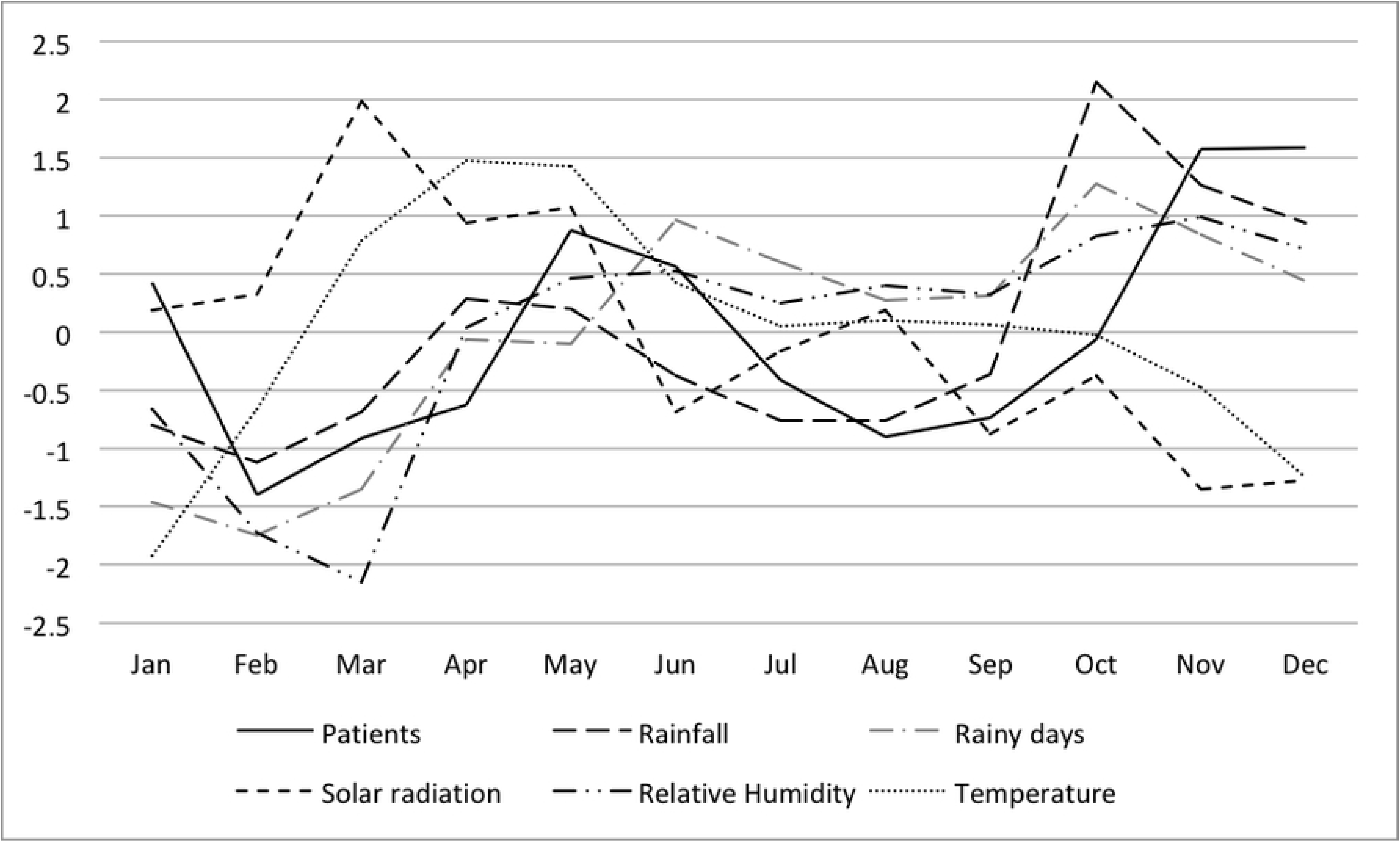
Monthly distribution of the standardized incidence of leptospirosis cases and meteorological parameters in the highlands of Sri Lanka.

Rainfall, the number of rainy days and relative humidity showed fluctuations similar to those of patient numbers, whereas solar radiation and temperature showed opposite fluctuations with respect to patient numbers for all three zones. In the dry zone, relative humidity fluctuated in a very similar pattern to the number of patients. Rainfall, the number of rainy days and relative humidity peaks were observed before the patient peaks in the highlands and dry zone; however, this relationship was not OR consistent] in the wet zone.

### Correlation analysis between variables

Before proceeding to the time series analysis, we analysed the association between the average number of patients and meteorological parameters using the Spearman correlation coefficient (Table 4). Both rainfall and the number of rainy days showed a statistically significant positive correlation in the dry and wet zones, but the associations were not significant in the highlands. Relative humidity showed a significant positive correlation across all three zones, whereas temperature showed a significant negative correlation across all three zones.

**Table 4:** Correlation coefficient (*r*) between the number of patients with leptospirosis and individual meteorological variables for each of the climatic zones.

| Variable | Dry Zone |  | Wet Zone |  | High Lands |  |
| --- | --- | --- | --- | --- | --- | --- |
|  | r value | p value | r value | p value | r value | p value |
| Rain fall | +0.21 | 0.010 | +0.31 | <0.000 | +0.09 | 0.265 |
| Rainy days per month | +0.22 | 0.005 | +0.23 | 0.004 | +0.15 | 0.069 |
| Temperature | - 0.37 | <0.000 | -0.18 | 0.025 | - 0.25 | 0.002 |
| Relative Humidity | +0.40 | <0.000 | +0.26 | 0.001 | +0.19 | 0.020 |

**\**r* – Spearman correlation**

### Models using the previously reported leptospirosis (univariate models) for climate zones

SARIMA models were developed to fit the monthly fluctuations in leptospirosis for each zone. Fig 6 shows the distribution of monthly leptospirosis cases from January 2007 to April 2019 for the three geographical zones, and seasonality was clearly visible in this graph. The highest number of cases was reported from the wet zone, whereas the lowest was from the highlands.

**Fig 6.**
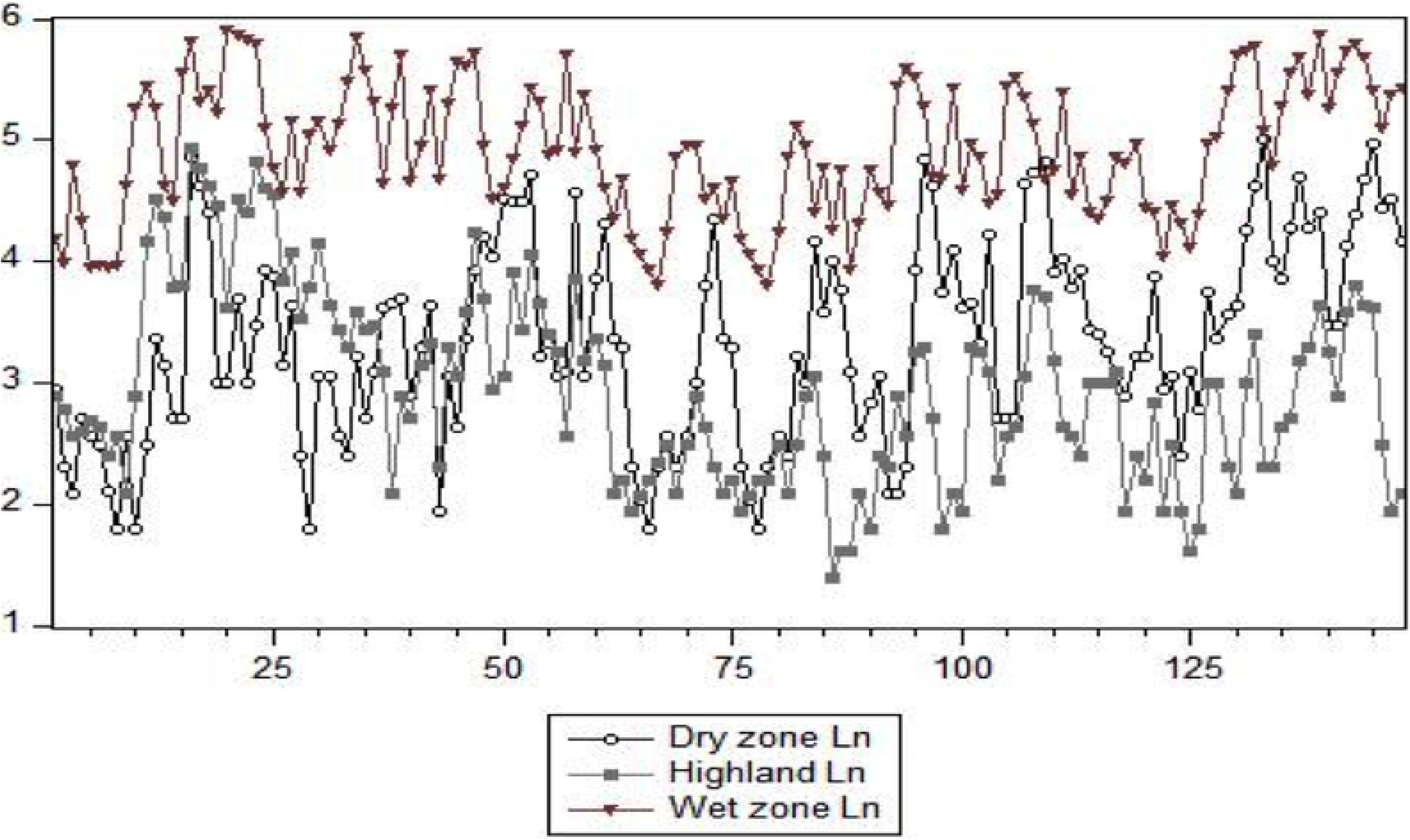
Monthly reported cases of leptospirosis from January 2007 to April 2019 (natural log—transformed data, Ln), by meteorological zones in Sri Lanka.

The best-fitting SARIMA models differed for the three zones, indicating different disease patterns (Table 5). All three models include substantial seasonal components, confirming the seasonality of leptospirosis.

**Table 5.** Best-fitting univariate SARIMA models to describe the incidence of leptospirosis in three climate zones in Sri Lanka.

| Zone | Model | Ljung-Box Q | Stationary R<sup>2</sup> | BIC | MAPE | Residual Normality |
| --- | --- | --- | --- | --- | --- | --- |
| Dry | ARIMA (1,0,0)(0,1,1)<sub>12</sub> | 0.23 | 0.71 | -1.0 | 12.81 | 0.65 |
| Wet | ARIMA (1,0,0)(1,1,1)<sub>12</sub> | 0.59 | 0.60 | -2.0 | 5.35 | 0.08 |
| High Land | ARIMA (0,1,1)(0,1,1)<sub>12</sub> | 0.53 | 0.59 | -1.2 | 14.29 | 0.84 |

**Table 6.** Unit root test for variables to assess the order of integration.

| Zone | Variable | Level t | Level p | 1<sup>st</sup> diff. t | 1<sup>st</sup>diff.p |
| --- | --- | --- | --- | --- | --- |
| Dry | Patient number | -5.77 | 0.0000 |  |  |
|  | Rain fall | -7.62 | 0.0000 |  |  |
|  | Rainy days | -2.79 | 0.0609 | -11.86 | 0.0000 |
|  | Temperature | -2.47 | 0.1226 | -12.91 | 0.0000 |
|  | Relative Humidity | -1.96 | 0.3010 | -12.52 | 0.0000 |
| Wet | Patient number | -5.07 | 0.0000 |  |  |
|  | Rain fall | -11.07 | 0.0000 |  |  |
|  | Rainy days | -8.80 | 0.0000 |  |  |
|  | Temperature | -2.15 | 0.2243 | -5.65 | 0.0000 |
|  | Relative Humidity | -2.40 | 0.1436 | -10.24 | 0.0000 |
| Hill country | Patient number | -4.15 | 0.0011 |  |  |
|  | Rain fall | -9.06 | 0.0000 |  |  |
|  | Rainy days | -8.04 | 0.0000 |  |  |
|  | Temperature | -3.69 | 0.0052 |  |  |
|  | Relative Humidity | -8.21 | 0.0000 |  |  |

\* t = t value ; p = p value

### Models using climate data and leptospirosis data (multivariable analysis)

We used the unit root test to determine whether the variable means were stationary (Table 6). The order of integration of the variables was not consistent for the dry and wet zones, and the mean was stationary for all variables for the highlands. Therefore, the ARDL model was used for further analysis.

Table 7 summarizes the results of the multivariable analysis. In the dry zone, the lag 1 effect of the dependent variable and the lag 1 and 2 effect of monthly rainy days showed significant association with the number of patients in the current month. None of the other variables were significant for the dry zone. In the wet zone, lag 1 and 3 of the dependent variable were positively associated. Lag 5 of rainfall was positively associated, whereas lag 2 and 3 were negatively associated. Lag 2 and 3 of monthly rainy days were positively associated, and this association was statistically stronger than the effect of rainfall. Lag 0 of monthly average temperature had a positive association, whereas lag 1 had a negative association. In the highlands, lag 1 and 2 of the dependent variable showed a positive association. Also, lag 1 of rainy days and lag 0 of temperature showed positive associations with the incidence of leptospirosis. For all three zones, relative humidity was not significantly associated with disease incidence, which is in contrast to findings from studies conducted in other parts of the world(11).

**Table 7.**
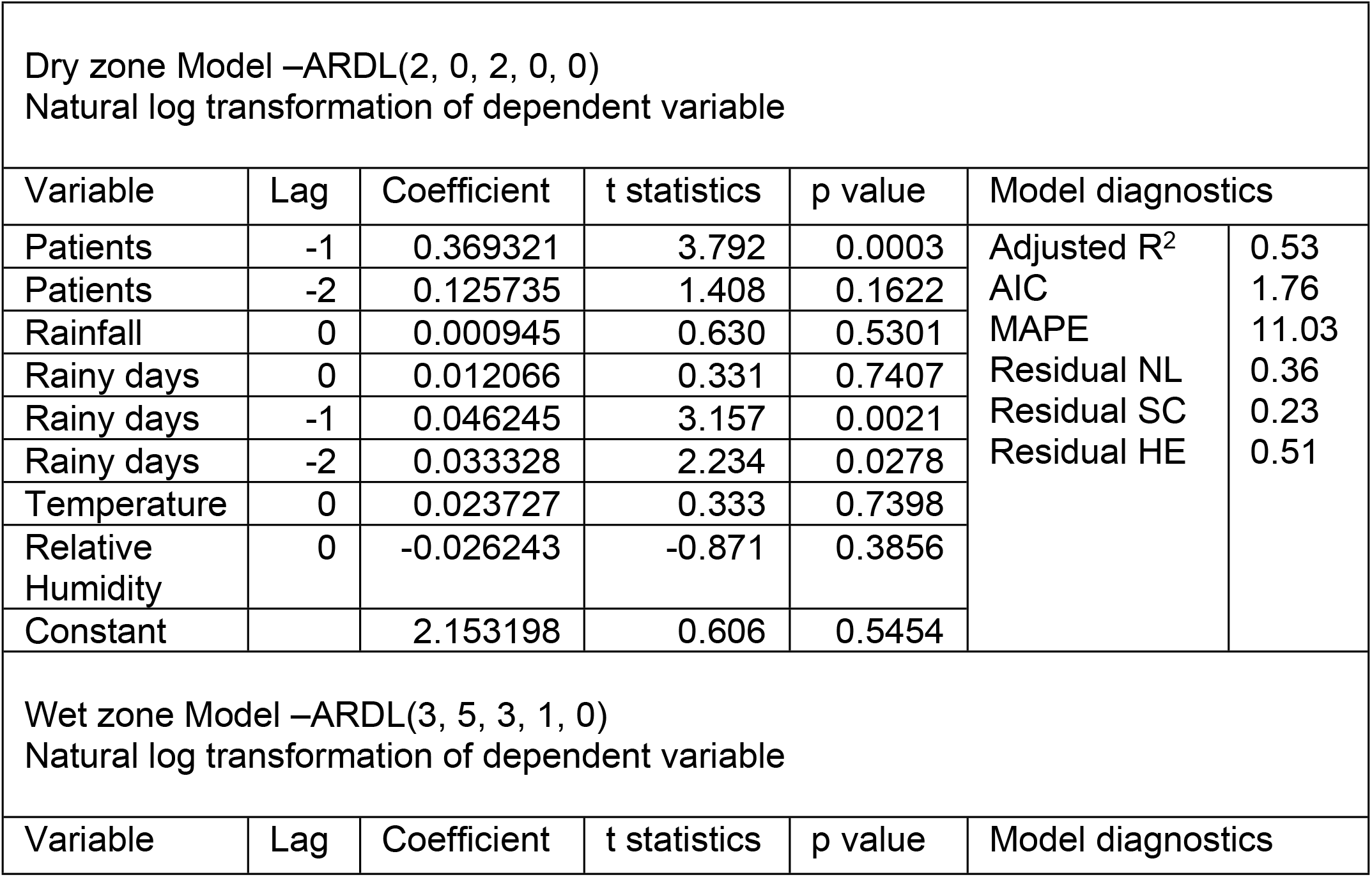
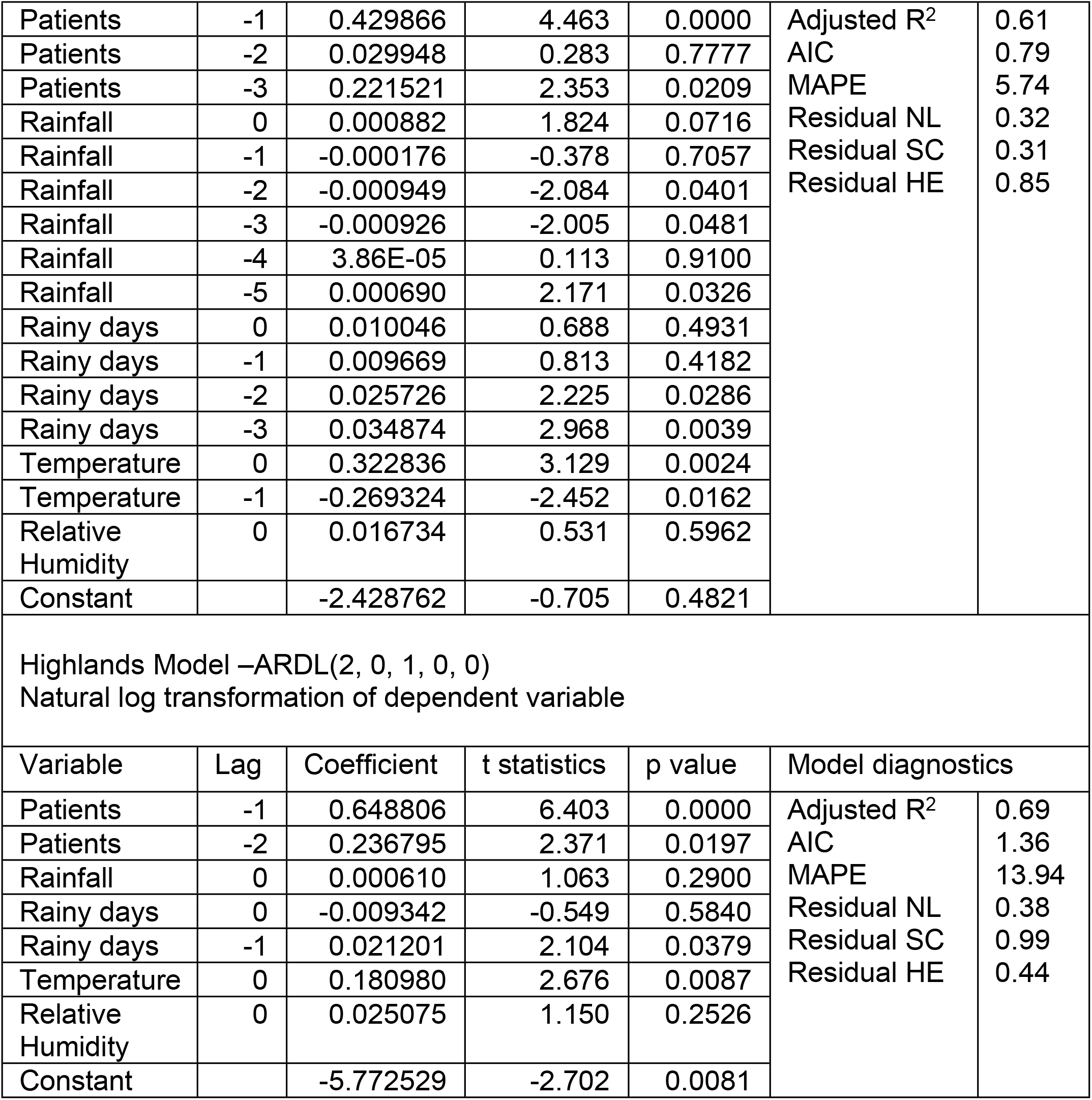
Multivariable ARDL models for the three climate zones.

Fig 7 shows the natural log—transformed observed and predicted values of leptospirosis cases for the three geographical zones.

**Fig 7.**
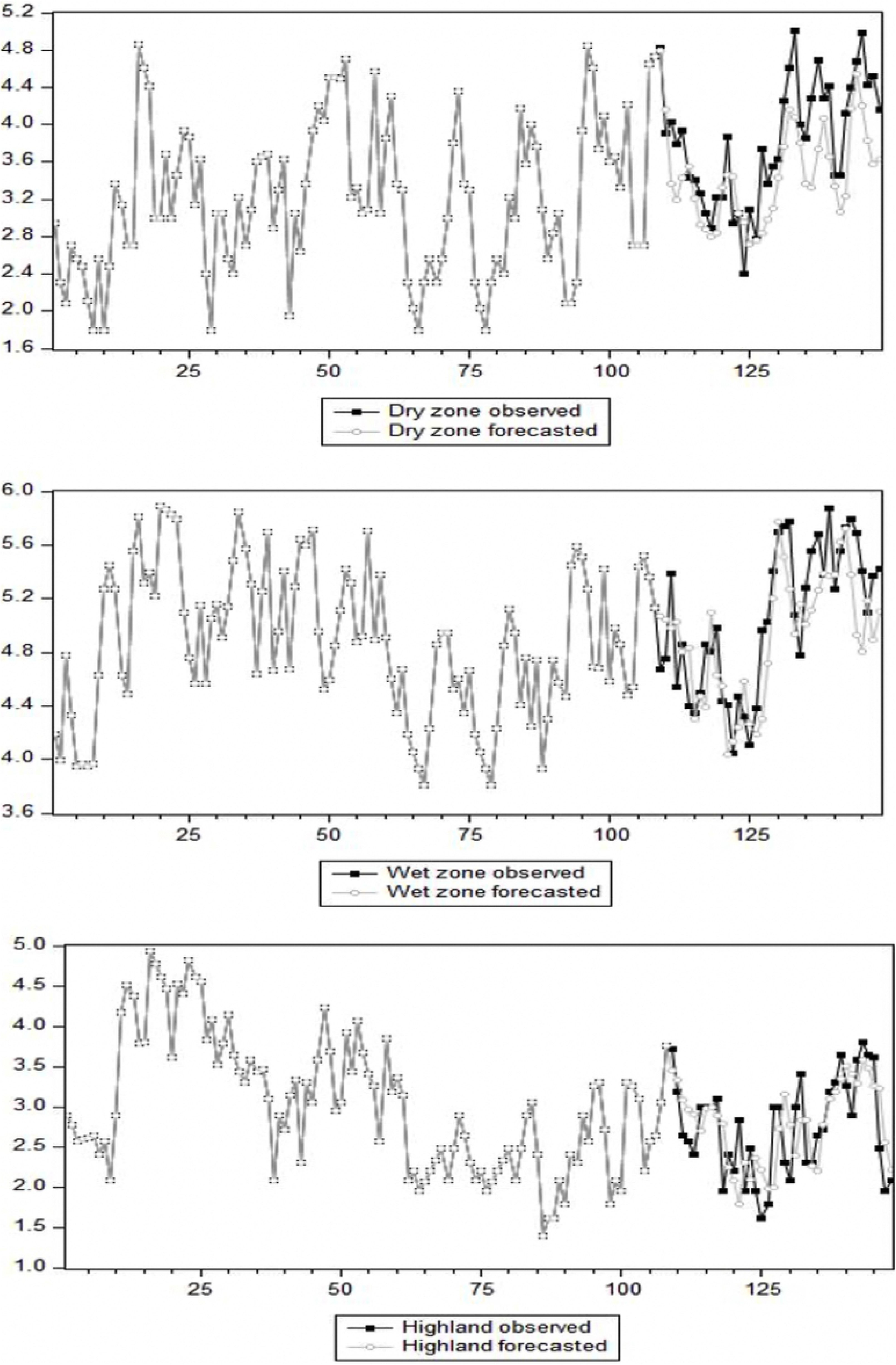
Natural log—transformed observed and forecasted values of the leptospirosis cases for the three climate zones.

## Discussion

Infectious disease modelling using administrative boundaries has been done in Sri Lanka previously with a focus on one or few districts. Leptospirosis, one of the two leading communicable diseases in Sri Lanka has never been modeled to understand the true effect of climate zones on disease incidence. In this study, we carried out a factor analysis based on rainfall data to confirm the climate zone and a time series analysis of leptospirosis disease incidence was done by climate zones.

As expected, the highest incidence of leptospirosis was reported for the wet zone, which also had the highest rainfall and highest number of rainy days per month. Frequent and sustained wetness of the soil due to heavy rainfall and frequent flooding may have contributed to the high number of patients in the wet zone(9,11). This finding was not, however, consistent across the dry zone and highlands. Although higher rainfall and a greater number of rainy days per month were reported for the highlands relative to the dry zone, the incidence of leptospirosis was not statistically different between these two zones. Lower temperatures and high solar radiation may have had a negative effect on the incidence of leptospirosis in the highlands. Another possibility is that the soil was not retaining water due steep slope in the highlands as compared with the lower-elevation dry zone, making it difficult for *Leptospira* to survive for a longer period. The survival of *Leptospira* in different climate zones has not yet been thoroughly studied, which makes further explanation of these findings difficult. A recently published study revealed that virulence of *Leptospira interrogans* is maintained for more than one year, even with prolonged starvation.(33) This observation provides an explanation for continuous outbreaks occurring in the wet zone. It also underscores the need for scientific studies on the natural survival of *Leptospira* under different environmental conditions.

Monthly fluctuation of rainfall, rainy days and relative humidity are suggestive of positive associations with leptospirosis (Figs 3—5). In the wet zone, however, the association at the early part of the year was different from that of the latter part of the year. Peaks in relative humidity, rainfall and rainy days were observed after the patient peak during the early outbreak, and peaks were observed almost simultaneously with the outbreak during the latter part of the year. Although there is no exact explanation for this variation, it might be associated with human behaviour associated with rice paddy cultivation. Paddy cultivation is the main occupational risk factor for leptospirosis in Sri Lanka(19). There are two main paddy cultivation seasons in Sri Lanka, referred to as ‘Yala’ and ‘Maha’. The Yala and Maha seasons differ slightly between the wet and dry zones. In the wet zone, paddy land preparation for cultivation for Yala starts at the end of February, and farmers are in regular contact with the mud during March, which is not part of the rainy season in the wet zone. This may explain the isolated patient peak observed during March in the wet zone. The rainy season from May to September and paddy land preparation for Maha from September onwards represent a possible explanation for the second patient peak in the wet zone. This may explain why rainfall has little effect on disease incidence, as the area is wet throughout the year in the wet zone.

Because outbreaks occur in specific climate zones, zonal-level models are of vital importance for predicting leptospirosis. The fitted univariate models differed for the three zones, providing additional support for different disease patterns. The seasonal component of the SARIMA models was significant for all three zones, consistent with a study in Thailand that found different disease patterns in Northern Thailand and Northeastern Thailand(13).

A correlation analysis together with multivariable ARDL models provided additional explanation for the meteorological associations. The average number of rainy days per month showed a consistent positive association with patient numbers for all zones. This association was consistent across several lag effects. Continues rain over a period of a few months may create a favourable environment for the growth and survival of *Leptospira.* Although several Sri Lankan and outside studies suggest that rainfall is a strong factor in predicting leptospirosis incidence, the average number of rainy days was highly associated variable than rainfall in this study. Especially in the highlands, rainfall was not correlated with the incidence of leptospirosis. This could be due to the geography of the highlands, as the mountains result in less water retention. Also, marshlands and flooding are not common in the highlands. Even in the multivariable analysis, rainfall was significantly associated with leptospirosis incidence only for the wet zone at lag 0. In contrast, rainfall showed a negative association at lag 2, 3 and 5 for the wet zone. There are studies suggesting that only limited rainfall is associated with vector-borne diseases(16,34). One study conducted in China indicated that heavy rainfall is negatively associated with leptospirosis when the natural habitats of rodents are destroyed(34). This suggests that light, steady rain over several days might promote leptospirosis more than a single heavy rainfall In contrast, a study in the United States found that floods can wash *L.interrogans* into surface water and spread the disease(35). Also, a Sri Lankan study conducted to compare two outbreaks revealed that *L. interrogans* was the predominant *Leptospira* species in the wet zone during the 2008 outbreak, whereas *L. kirschneri* was the predominant *Leptospira* species in a post-flooding outbreak of leptospirosis in the dry zone (31). Floods are more common in the wet zone than in other zones. The link between *L. interrogans* and flooding may explain the isolated significance of rainfall in the wet zone. This may also explain the lack of significance for rainfall in the highlands.

Although the correlation between relative humidity and the number of patients was significant, the effect was not significant in the multivariable analysis. This may be due to the presence of multicollinearity in the correlation study, although the variance inflation factor for relative humidity was <5. There are several studies suggesting that warm, humid conditions are needed for survival of *L. interrogans* outside the host(12,15). Although some studies suggest that a higher relative humidity has a negative impact on rodents(36,37), there is evidence suggesting that the primary host of *Leptospira* in Sri Lanka may not be rodents(6,38). Thus we need further studies on the possible hosts of *Leptospira* in Sri Lanka.

The average temperature showed different associations with disease incidence across the three zones. As the dry zone has a higher temperature throughout the year, there may not be an additional effect of temperature on leptospirosis in this zone. The average temperature in the highlands is below the optimal temperature for the growth and survival of *Leptospira.* This may explain the positive association of temperature in the highlands. In the wet zone, temperature showed a positive effect at lag 0 and a negative effect at lag 1. Although an exact reason cannot be given for these contrasting effects, temperature is a positive factor associated with the survival of rodents and rodent densities(15). Further studies are needed to understand the association of temperature with leptospirosis.

Solar radiation was not included in the multivariable model because of missing values in the data set. However, the correlation analysis did show that solar radiation was negatively associated with the number of patients in the dry zone and highlands, whereas there was no association in the wet zone. Bacteria grow well under conditions of high solar radiation, whereas solar UV may inhibit the growth of bacteria. *L. interrogans* can survive for up to 6 hours under UV exposure(39). There is also evidence that bacterial growth is optimal during the rainy season because of lower doses of radiation(40). However, this evidence does not provide adequate explanation for the observations in this study.

This study aimed to provide a meteorological explanation for the variability of leptospirosis across different geographical zones in Sri Lanka. This variability includes differences in clinical features, unusual clinical manifestations and differences in the severity of outbreaks that have occurred since the major outbreak of 2008(18). However, a meteorological explanation alone is not sufficient to answer the question. We also need to consider factors that have direct effects on human behaviours, host behaviours, host reproduction, susceptibility and bacterial growth and survival. Studies on host animals, as well as on the geographical distribution of etiological agents, the microbiome and microbial survival in different soil types and water sources are crucial for describing the diversity of leptospirosis and to obtain a holistic understanding of the disease patterns(41).

## Limitations of the study

First, leptospirosis is underestimated in Sri Lanka(29) which is a common scenario globally due to the nature of the disease. Lack of clinical suspicion, unspecific presentations and lack of point of care diagnostics has been showed to be contributing to this. Yet, the above mentioned underestimations are everywhere in the country thus the effect of that possible underestimation might be having a minimal effect.

## Abbreviations

AIC: Akaike information criterion
AR: Auto Regressive
ARCH: Auto Regressive conditional heteroscedasticity
ARDL: Auto Regressive distributed lag
ARIMA: Auto regressive integrated moving average
ARIMAX: Multivariate auto regressive integrated moving average with input series
DALY: Disability adjusted life years
IQR: Inter quartile range
LM: Lagrange multiplier
MA: Moving Average
MAPE: Mean absolute percentage error
SARIMA: Seasonal Auto regressive integrated moving average with input series
SPSS: Statistical package for social sciences
UV: Ultraviolet
VAR: Vector Auto Regressive

## Acknowledgements

We acknowledge the epidemiological unit of Sri Lanka for giving permission to use the published leptospirosis data from the weekly epidemiological reports. Also, we acknowledge the medical students, and the staff of faculty of Medicine and Allied Sciences, Rajarata University of Sri Lanka for the support given for data entering. We also thank Mr. Dulara Piyumal for his help with generating the figures.

## Funding

Data from the meteorological department were purchased using the dean’s award for research publications received by JW in 2017.

## Availability of data and materials

All of the meteorological data are available from the meteorological department of Sri Lanka, and epidemiological data are freely available on the website of the epidemiology unit of Sri Lanka.

## Data availability statement

The data that support the findings of this study are available in [Zenodo] at http://doi.org/[10.5281/zenodo.4290803].

## Author contributions

JW was involved in the research design, literature search, data extraction, data interpretation and final report writing. SA conceived the study, guided the research process, reviewed the extracted data and reviewed and edited the final manuscript. RA guided for the research proposal, the research process and data analysis and edited the final manuscript.

## Ethics approval

This study was conducted using secondary data, and all the data used are publicly available. Therefore, ethics approval was not obtained.

## Consent for publication

Not applicable

## Competing interest

The authors declare that they have no competing interests.

